# Spatial behaviors and seasonal habitat use of the increasingly endangered thick-billed parrot (*Rhynchopsitta pachyrhyncha*)

**DOI:** 10.1101/2023.08.30.555620

**Authors:** James K Sheppard, Javier Cruz, Luz Francelia Torres González, Miguel Ángel Cruz Nieto, Ronald R. Swaisgood, Nadine Lamberski

## Abstract

The thick-billed parrot (*Rhynchopsitta pachyrhyncha*) inhabits highland pine forests in the Sierra Madre Occidental ranges of northwestern Mexico. Their populations have declined significantly to <2000 individuals due to habitat loss, illegal hunting and increasing predation. Despite their ecological and cultural importance and increasingly endangered status, the species is data deficient. Our study aimed to inform and enhance conservation management strategies for thick-billed parrots with information on their spatial ecology, habitat use, migratory behaviors and social associations. We deployed biotelemetry devices to conduct the first tracking study of wild thick-billed parrots. Our study revealed that thick-billed parrots are seasonal migrators, departing their breeding habitats around October and returning from southern habitats around April. Our research also identified previously unknown overwintering sites and migratory stopover locations, as well as a new nesting site. The parrots exhibited high spatial variability in range shifting behavior, but all tracked parrots exhibited range shifts during migration, with durations of 3 to 181 days and distances of 173 to 765 km. They traveled in close social groups and migratory routes primarily followed high-elevation forests along the Sierra Madre Occidental ranges. Home range analysis indicated smaller breeding site ranges and larger overwintering ranges, possibly reflecting nesting constraints and winter food resource dispersion. Parrot spatial associations favored high-elevation forest landscapes with tall and wide-trunked trees, underscoring the importance of preserving old-growth forests for nesting and foraging. Less than 20 % of parrot habitats have formal regulatory protections. Conservation management efforts must focus on increasing protections for nesting areas, overwintering habitats, and key migratory stopover sites. As climate change exacerbates regional threats, integrated management plans involving local stakeholders and communities are essential for the parrots’ long-term survival and the preservation of their old-growth forest habitats.

## Introduction

Thick-billed parrots (*Rhynchopsitta pachyrhyncha*) inhabit highland pine forests at elevations above 2000 m in the Sierra Madre Occidental ranges of northwestern Mexico where their breeding ranges are located. The parrots also overwinter within the south-central Sierra Madre Occidental. The species is highly mobile and historically ranged across the US-Mexico border into southeastern Arizona and possibly southern New Mexico, where it has been extirpated since 1938 likely due to excessive unregulated shooting (Lanning, Dirk V and Shiflett, James T 1983, Snyder et al. 1994, 1995, Cruz-Nieto, M.A. 1998, Monterrubio et al. 2002, Monterrubio-Rico et al. 2015). The parrots nest in tree cavities and feed primarily on seeds from several high-elevation pine species, including *Pinus strobiformis, P. arizonica*, and *P. durangensis* (Lanning, Dirk V and Shiflett, James T 1983, Snyder et al. 1994).

The Mexican government and the U.S. Fish and Wildlife Service currently list the thick-billed parrot as ‘Endangered’ (U.S. Fish and Wildlife Service. 2013). The species is listed in Appendix I of CITES, and the International Union for Conservation of Nature also lists its status at ‘Endangered’ because of its decreasing populations, specialized conifer diet, limited geographic range, and niche as a cavity-nesting bird (IUCN 2017). Historical population declines occurred in Mexico from unregulated pet trading and more recently from increased logging within the parrots’ range (Lanning and Shiflett 1983). Availability of nesting cavities in mature trees and snags has dramatically decreased in recent decades due to destructive clear-cutting and high-intensity timbering of remnant old-growth primary forests, as well as higher incidences of catastrophic wildfires exacerbated by drier conditions due to climate change (Sáenz-Romero et al. 2010, Monterrubio-Rico et al. 2015). Surveys of thick-billed parrot populations conducted over the past 25 years within their remaining breeding sites show a decline of 20 – 30 %. The most recent surveys indicate there may be fewer than 2000 individual birds extant (Sheppard et al. 2021), although this number is questionable due to the irregularity of survey intensity.

This precipitous drop in thick-billed parrot populations has been linked to the decline in suitable nesting resources (Monterrubio-Rico and Enkerlin-Hoeflich 2004), as well as a trend observed in recent years of increasing predation events on nesting thick-billed parrots and their chicks (Sheppard et al. 2021). Less than 20 % of the thick-billed parrot population will typically breed annually, so the species can persist and recover only if sufficient nesting resources are available (Monterrubio et al. 2002). Unfortunately, less than 1 % of the old-growth forest that once covered Mexico’s Sierra Madre Occidental remains (Lanning et al. 1983). Consequently, a revision of the population status of the species to ‘Critically Endangered’ status may be warranted, based on criteria A.2. of the IUCN Red List: ‘An observed, estimated, inferred or suspected population size reduction of 80% over the last 10 years or three generations, whichever is the longer, where the reduction or its causes may not have ceased OR may not be understood OR may not be reversible’ (IUCN 2012).

Despite their iconic profile, ecological and cultural importance, and increasingly threatened populations, thick-billed parrots are highly data-deficient. Basic information on resource requirements is lacking for the vast majority of Psittaciformes (Renton et al. 2015, Berkunsky et al. 2017), and the few formal studies describing thick-billed parrot habitat and resource use were published in the 1990s. There are no empirical investigations of the spatiotemporal characteristics of thick-billed parrot home ranges or their migratory paths, and their phenological responses to seasonality remain unelucidated. While the locations of thick-billed parrot nesting sites in the northern reaches of their range are known and have been studied for decades by field monitoring and *in situ* health management programs, the locations of their southern overwintering sites remain largely unknown and undefined. Hence, there is an urgent need for conservation management strategies for thick-billed parrots and their high-elevation forest habitats to be informed by updated information on their spatial behaviors and ecological requirements. Specifically, conservation managers require information on where and when these parrots spend their time, and the preferred characteristics of habitats along the migratory routes, in order to devise targeted conservation interventions.

Advances in the power and miniaturization of wildlife biotelemetry devices in tandem with new empirical methods to model large and complex spatial ecology datasets are enabling previously unobtainable insights to be obtained on the spatial behaviors of an expanding range of avian species (Bridge et al. 2011, Kays et al. 2015, Williams et al. 2020). We capitalized on recent reductions in the size and weight of solar-powered Argos transmitters to conduct the first remote tracking study of wild thick-billed parrots. Our overarching goals (paraphrased from the current thick-billed parrot recovery plan, section 3.0. pages 58+: U.S. Fish and Wildlife Service. 2013) are to help guide the development of a long-term habitat conservation plan for the evaluation of habitat availability and use for thick-billed parrots and promote habitat restoration for the species within their foraging areas and nesting sites. To meet these goals, our objectives were to collect detailed information on: 1) How these birds choose and use their habitats; 2) how they move between their breeding and overwintering ranges; 3) the locations and dimensions of thick-billed parrot southern overwintering ranges and; 4) how the parrots spatiotemporally associate amongst themselves across seasons.

Previous studies and anecdotal evidence indicate that thick-billed parrots spend their breeding season (June to October) primarily within the states of Chihuahua and northern Durango, then overwinter (November to May) within the south-central Sierra Madre Occidental in the states of Durango, Jalisco, Michoacán, Nayarit, and Colima (Snyder et al. 1994, Cruz-Nieto, M.A. 1998, Monterrubio et al. 2002, Monterrubio-Rico et al. 2015). However, there are no published accounts of the locations, configuration, or dimensions of these overwintering ranges, nor have the seasonal migratory routes used by thick-billed parrots or the presence of their migratory stopover sites been formally investigated and described. We predicted that the parrots leave breeding areas at a certain time each year and return at another time, and the timing, directions and distances of their migrations are relatively constant and predictable from year to year. We also predicted they depart their seasonal ranges in social groups within four to six weeks and consistently travel in flocks along similar routes when migrating either north or south. To test these predictions, we deployed transmitters on a representative sample of free-ranging thick-billed parrots and tracked their movements and range use patterns across multiple seasons.

## Methods

### Study area and biotelemetry

All thick-billed parrots were captured and telemetered during the breeding season (September – October) at nesting sites within the Sierra Madre Occidental ranges of northwestern Mexico, State of Chihuahua from 2019 to 2022. These ranges are the longest in Mexico, with a rugged physiography of highland plateaus and deep canyons encompassing elevations from 300 to 3300 m. The Sierra Madre Occidental extends more than 1160 km, from the Madrean Sky Islands of southern Arizona and New Mexico (30° 35’N) to northern Jalisco (21°00’N) in western Mexico and is the source of important environmental services for many communities and industries in northwestern and north-central Mexico. The Sierra Madre Occidental forest ecoregions also support some of the greatest biodiversity in North America (González-Elizondo et al. 2013), and comprise an important biological corridor for plants and animals (Kobelkowsky-Vidrio et al. 2014).

We deployed 9.5g solar-powered PTT transmitters (Microwave Telemetry Inc.) on each tracked parrot using a custom-fitted backpack harness constructed of Spectra™ material (Balley Mills Inc.) (Fig.3). Transmitters were 2.6 % of the mean 361.7 g (± 4.5 SE) bodyweight of the captured birds (Appendix 1). Adult and fledgling parrots were retrieved from nesting hollows or artificial nesting boxes located on the trunks of pine trees (Arizona pine, Apache pine, and Chihuahua pine) about 6 – 12 m from the ground. Health checks and morphometric data were collected from each parrot prior to attachment of the transmitter and harness, with the bird returned to its nest after a total handling period of < 20 minutes. Of the 34 parrots telemetered, 20 were breeding adults and 14 were fledglings with an average estimated age of 55 days (Appendix 1).

### Data post-processing

Transmitter location data were calculated by the CLS-ARGOS service using their Kalman filter algorithm (Lopez et al. 2014). All subsequent data processing and statistical analyses were performed using ‘R’ language (version 3.6.1, R Core Team 2019), with mapping of location data and geospatial data processing conducted in ArcGIS Pro (ESRI Inc). To account for location uncertainty and the irregular time series of Argos positions, we used the continuous-time, correlated random walk (CRW) state−space model in the R package *aniMotum* (Jonsen et al. 2019, 2020, 2022). We used the CRW model to estimate time-regularized (12 h) intervals that predict the most probable paths flown by each bird. Large gaps in tracking data can be problematic when fitting CRW models, as the lack of data may lead to implausible predictions in these gaps. Therefore, prior to CRW modeling and statistical analysis, tracking data with a gap (here defined as ∼18 h or longer) were trimmed to remove the gap.

### Data analysis

We used the time-varying move persistence model in the *aniMotum* package to objectively identify changes in the filtered and regularized thick-billed parrot movement trajectories,

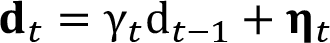

where displacements **d**_*t*_ = ***x***_*t*_ − ***x***_*t*−1_ and **d**_*t*−1_ = ***x***_*t*−1_ − ***x***_*t*−2_ are the changes in an animal’s location ***x*** at times *t* and *t* − 1. The random variable **η**_*t*_ = N(**0, Σ**), with variance–covariance matrix **Σ** specifying the magnitude of variability in the two-dimensional movements. *γ*_*t*_ is the time-varying move persistence between displacements **d**_*t*_ and **d**_*t*−1_ (Jonsen et al. 2019). *γ*_*t*_ is an index of behavior that is continuous-valued between 0 (low move persistence) and 1 (high move persistence). We imported the filtered and regularized movement data into the *marcher* package for R to conduct the mechanistic range shift analysis method of Gurarie et al. (2017). This method can account for the autocorrelation and irregular sampling of telemetry movement datasets to enable quantification of the extent of a range shifts, as well as tests for stopovers and site fidelity tests. We generated likelihood estimates for hypothesis testing of range shifts and stop-overs during parrot migrations and performed the range shift tests against a null hypothesis of no range shift. We compared fitted models with and without a single range shift with a likelihood ratio test (l.r.t. with 4 d.f) corresponding to the estimation of two more range center coordinates and times of migration. Range shifting refers to the movement of species or populations to a new geographic area in response to changing environmental conditions (Parmesan 2006). We used the range shift index (RSI) method (Gurarie et al. 2017) as a continuous index of the relative extent of a parrot’s range shift, defined as the ratio of the distance from the x, y locations of the range shift centroids to the diameter of the ranging area. An RSI of 0 indicates no range shift and increasing values indicate longer migrations. The *P*-value of the range test quantified the ‘detectability’ of a range shift, while the RSI quantified the ‘effect size’. Many bird species alternate long-distance movements with time spent at a stopover – the time and space where individuals rest and refuel for a subsequent migratory flight (Taylor et al. 2011). Stopover sites can have a large influence on the rate of mass gain in migrating birds, which can have consequences for the timing and success of reproduction, as well as survival during the stationary non-breeding season (Sheehy et al. 2011, and references within). Hence, we also conducted a stopover test for each parrot by fitting a model with three ranges against a null model of two ranges to identify any stations within a migration (l.r.t. with 4 d.f.). Finally, for parrots that exhibited consecutive seasonal range shifts, we analyzed their migratory paths separately as either orientations towards the south away from the nesting sites or northerly return migrations.

We summarized the results by comparing the occurrence of range shifting (determined by the α = 0·05 significance test), and exploring the relationships between seasonal movement direction, duration, and distance and ranging area following Gurarie et al. (2017). Post hoc analyses were performed with linear mixed effects models with individual parrots as random effects (using the *lmerTest* package in R; (Kuznetsova et al. 2017)). We used a Kruskal-Wallis one-way ANOVA in the *ggstatsplot* package for R to explore differences in parrot dispersal patterns across the entire tracking period (Patil 2021) and to test whether parrots tracked within month groups had the same distribution of scores for each tracking year using median net squared displacement as the quantitative response variable. We used a Dunn post-hoc test to indicate the greatest pairwise difference in NSD between months. We also used the R package *nestR* to detect the presence of previously unknown nesting sites by identifying temporal persistence of recursive movements made by individual tracked parrots to locations within the northern breeding habitats during the breeding season. The *nestR* method identifies repeatedly visited locations along movement trajectories, with the inference that these patterns enable determination that a revisited location is likely to be a nest (Picardi et al. 2020).

We used the integrated step-selection function in the *amt* package for R to fit a simple Habitat-Selection Function (HSF) model to the parrot movement data (Signer et al. 2019, Fieberg et al. 2021). The HSF connects logistic regression, Inhomogeneous Poisson Point-Process (IPP) models, and weighted distribution theory to estimate the probability of a particular habitat being used disproportionally relative to its availability compared to a reference category. Positive model coefficients covariate effects (β) for a given class variable indicate a higher likelihood of use (i.e., selection) compared to the reference class, while negative coefficients indicate a lower likelihood of use (i.e., avoidance) (Fieberg et al. 2021). We examined five potential environmental correlates of thick-billed parrot spatial association: Elevation of the underlying terrain, average tree height, tree trunk diameter, tree density, and forest species, using elevation as the reference category. Terrain elevation was derived using the Advanced Spaceborne Thermal Emission and Reflection Radiometer (ASTER DEM V2) 1 arc-second, 30 m resolution elevation grid dataset for Mexico (Tachikawa et al. 2011). Information on tree characteristics was sourced from the Open Data Portal, National Forest and Soil Inventory of the Mexican Government (INEGI 2021, CONAFOR 2023). We converted a vegetation geospatial layer from the INEGI portal into a continuous predictor raster with pixels assigned to the following rating scale of increasing habitat quality based on prior knowledge of parrot habitat association: 1 = grassland/shrub/agricultural, 2 = oak forest, 3 = mixed oak/conifer forest, 4 = conifer forest (Snyder et al. 1995, Monterrubio-Rico and Enkerlin-Hoeflich 2004).

Numerical covariate raster layers were scaled and appended to the state-space filtered parrot location data and a sample of random (‘available’) points was generated for each parrot in a weighted logistic regression model. The key assumptions for the IPP model were *a*) the total number of location points in the study region is a Poisson random variable, and *b*) that locations are independent (any clustering can be explained by spatial covariates). To meet these assumptions, we assigned a weight of 5000 to available locations and a weight of 1 to all used locations, with a significance level of α < 0.05 for the HSF variables (Signer et al. 2019, Fieberg et al. 2021). We fitted the HSF model to individual birds and treated the estimates as data (two-step approach), which provided a simple way to explore among-animal variability. We also used the *glmmTMB* package for R (Brooks et al. 2017) to generate an independent random slopes model and estimate population-level mean coefficients for each environmental covariate (mixed-model approach of Muff et al. 2020). We assumed that all landscape types were available to all tracked parrots. Additionally, we generated used-habitat calibration (UHC) plots to evaluate model performance by comparing the ‘used’ distributions of each habitat covariate in the model to its predicted distribution generated from the model (Appendix 3). If the model is well-calibrated, observed used habitat in the independent testing dataset will fall primarily within the confidence bounds of the predicted used distribution (Fieberg et al. 2018).

We fitted variograms and continuous-time movement models (CTMMs) to the filtered PTT location data using the *ctmm* package for R (Calabrese et al. 2016). We chose this continuous-time stochastic movement framework to model the parrot Argos data because: 1) It properly accounts for the serial autocorrelation intrinsic to movement data; 2) it explicitly accounts for telemetry error; 3) it is robust to small sample sizes; 4) it can handle irregularities and gaps in location datasets, and; 5) it appropriately estimates confidence intervals (for details on this method see (Calabrese et al. 2016, Fleming and Calabrese 2017, Papageorgiou et al. 2021, Silva et al. 2021). We used Autocorrelated Kernel Density Estimation (AKDE) methods to estimate individual and mean home range areas, which capture the long-term, 95% coverage region of the probability distribution of all possible locations given observed movement properties (Fleming et al. 2015). These models follow a continuous-time stochastic process from which the maximum likelihood Gaussian home range area can be extracted after the best fitted model is selected based on AIC. We generated separate AKDE home ranges for location data recorded at the northern breeding sites and the southern overwintering sites using the spatiotemporal delineation criteria identified from the mechanistic range shift analysis. We also used the outputs from the CTMM to calculate a mean Kriged occurrence distribution estimate of parrot movements during migrations between their two seasonal ranges to characterize population-level migratory route areas (using the 95% contour) and define the locations of stopover sites (using the 25% contour).

We used the Bhattacharyya coefficient, a metric initially derived to measure similarity among probability distributions (Bhattacharyya 1943), to examine spatial associations between parrots, and assign individuals to a social group when the Bhattacharyya coefficient overlap values > 0.5. This metric is proportional to areal overlap among distributions, does not depend on *ad hoc* parameters (such as isopleths), and incorporates a correction for small sample sizes (Winner et al. 2018, Mertes et al. 2019). We used the *ctmm* package to calculate bias-corrected Bhattacharyya coefficients and record confidence intervals for all pairwise comparisons of AKDE home ranges for each bird. Bhattacharyya coefficients range from 0 to 1, 1 if the two distributions are identical and 0 if there is no shared support. We characterized the long-term encounter location probabilities for parrot movements within their home ranges using the conditional distribution of encounters (CDE) function within the *ctmm* package (Noonan et al. 2021). We derived probability densities with confidence intervals using the CDE method to identify sites where group-level spatial interactions most likely occurred within the overlapping parrot home ranges. We also used a paired *t*-test to compare the AKDE home range areas and overlap values of parrots at nesting versus overwintering sites. Finally, we imported polygons of protected areas in northern Mexico acquired from the Mexican Government (INEGI 2021) into ArcGIS Pro. We then intersected these layers with the population-level polygons of thick-billed parrot home ranges to calculate the areas and percentages of parrot seasonal ranges that are currently covered by formal protections (e.g. habitat protection zoning).

## Results

Of the 34 parrots that were telemetered, 22 had enough filtered relocation fixes (>200) to conduct an analysis of spatial behaviors with sufficient rigor to model their movement behaviors and habitat selection. From these 22 parrots a total of 44 long-distance seasonal migration tracks were extracted (31 southern migrations, 13 northern migrations). Twelve parrots were tracked undertaking return migrations from the southern overwintering habitats back to the northern nesting sites, and six parrots were tracked across multi-year seasonal migrations. 74 % of southward migration events began in October as temperatures cooled and chicks produced in the breeding season fledged (16 % began earlier in September and 10 % began later in November). 77 % of northward migration events began in April as temperatures rose. A stopover was confirmed (*p* <0.01) in 16 of the 22 tracked parrots and in 23 of the 44 total migratory tracks (73% of parrots, 52% of tracks). Analyses of parrot trajectories revealed a high degree of variability in range shifting behavior among individuals and groups. Parameter estimates, confidence intervals and test statistics for the tracked parrots are summarized in Appendix 2. and presented visually in Figs. 2. and 3. All parrots had significant range shifts. Migration times were highly variable and ranged from 3 to 181 days (median 18, inter-quartile range 8-37). The distance of the shift ranged from 173 to 765 km (median 461, inter-quartile range 350-540), with the range shift index ranged from 1.0 to 12.3 (median 2.9, inter-quartile range 2.2-3.8).

**Fig 1.**
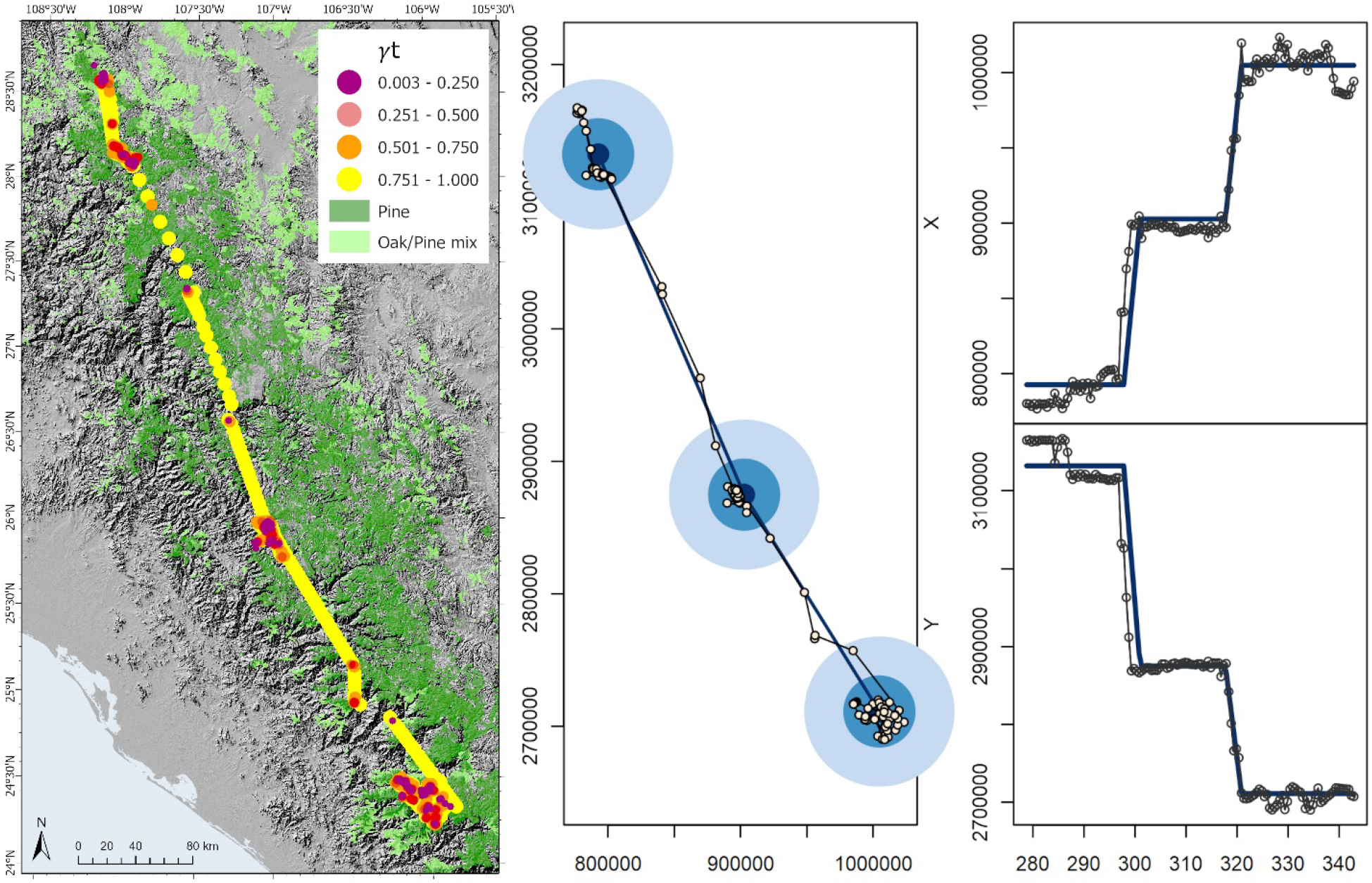
(Left) SSM-filtered thick-billed parrot tracks from a representative individual parrot (id = TBP23) telemetered at the Tutuaca nesting site in Oct-21 and tracked across a 500 km migration completed in 24 days from the northern nesting sites to the southern overwintering habitats. Location data were resampled at 12-hour intervals and colored according to the associated move persistence (*γ*_*t*_), continuous-valued between 0 (low move persistence) and 1 (high move persistence). Extent of primary growth high-elevation pine and pine-oak forest (green polygons) displayed for reference. (Center) Range shift plots circles are (dark to light) the 50% and 95% areas of use and indicate range shift plots with three ranges (two shifts) plotted on UTM easting (X axis) and northing (Y axis) coordinates in meters. (Right) Time series on the time series plot x-axis are in days from initial capture beginning 06-Oct with blue lines reflecting confidence intervals around the estimated means (also plotted against X-Y UTM coordinates). A range shift was confirmed by the model (d.f. = 4, l.r.t = 315, *p* <0.005, RSI = 4.0 (95% CI 3.8-4.3)), as was the stopover indicated by the central range circles and spatial shift in the middle of the time series (d.f. = 4, l.r.t = 113, *p* <0.005).

**Fig 2.**
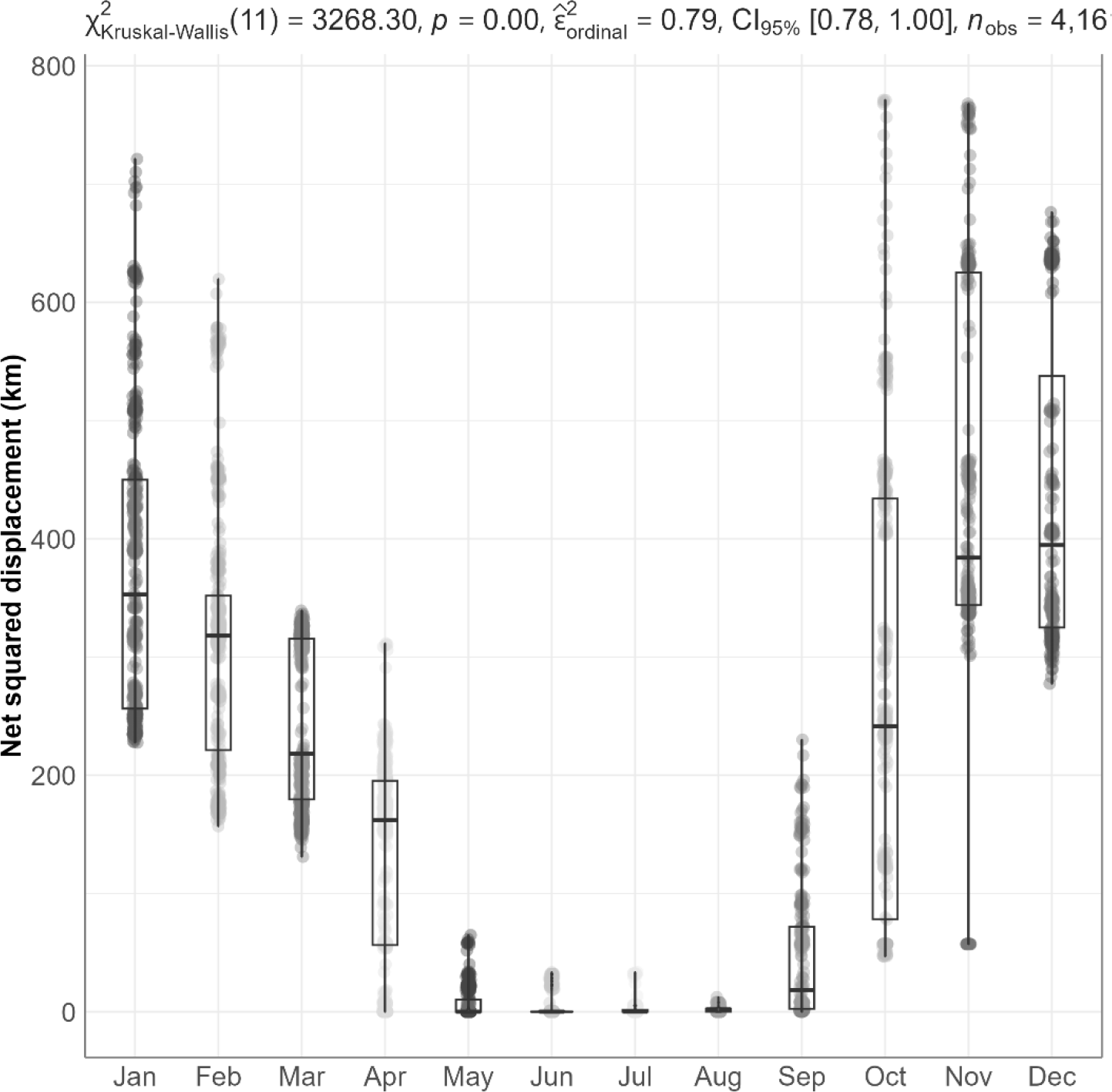
Monthly median net squared displacement (NSD) with IQR values in km for SSM-filtered thick-billed parrot tracks resampled at 12-hour intervals across tracking year 2022. A Dunn post-hoc test indicated the greatest pairwise difference in NSD occurred between January and July (p < 0.001).

**Fig 3.**
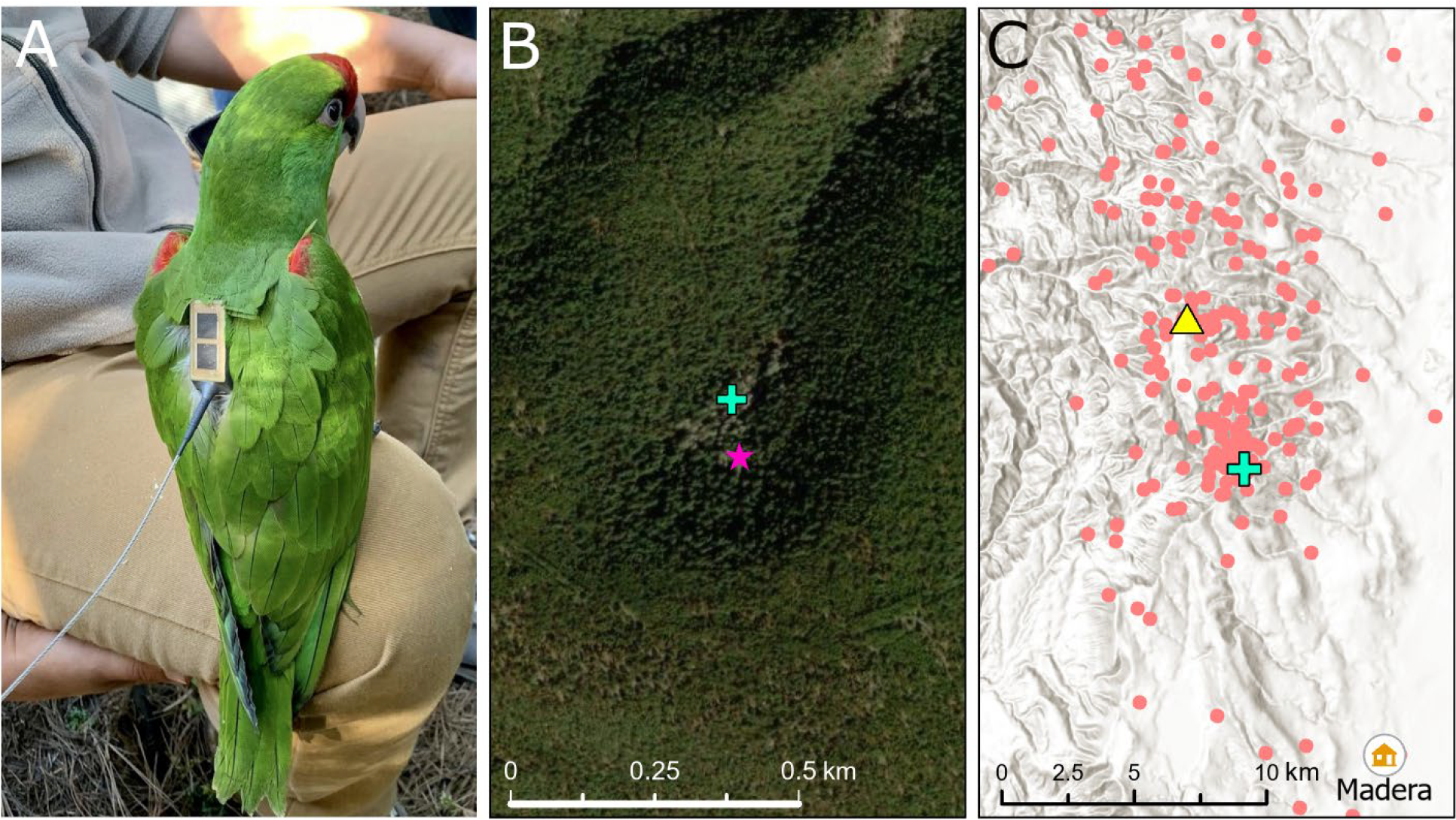
Location of a previously unrecorded 2022 parrot nest site near the Chihuahuan town of Madera that we predicted from the tracking data using the *nestR* package for R (Picardi et al. 2020). **A**. Backpack attachment method for the transmitter. **B**. Pink star indicates the predicted nest site location, and the blue cross indicates the actual nest location confirmed via ground-truthing. The zoomed-out topographic map on panel **C**. shows the spatial distribution of the tracking data (red dots) used to derive this nest site prediction, with the yellow triangle indicating the nest site where this bird was initially telemetered in 2021.

Migration duration or distance did not show any significant patterns with respect to whether the birds were migrating north or south (mixed effects model *p* > 0·6). However, there was a significant effect on migration duration if a stopover was included (*p* 0.01) which added an average of 22.6 days (± 8.8 SE) to the path. The tracked parrots spent an average of 146.7 days (± 10.6 SE) within their southern overwintering habitats before making the return migration north (Appendix 2.). Parrot net squared displacement distributions were significantly different among months, using Kruskal-Wallis (2020: X^2^ = (11) 869, *p* < 0.005. 2021: X^2^ = (11) 685, *p* < 0.005. 2022: X^2^ = (11) 1086, *p* < 0.005), and these scores were similar across tracking years 2020-2022. A Dunn post-hoc test indicated the greatest pairwise difference in NSD occurred between January and July (*p* < 0.001). NSD values began increasing in October when most parrots have begun migrating south, with a peak in December-January. NSD values then decreased to their lowest values in June-August, when the parrots had returned to their northern nesting grounds where their movements were concentrated within a localized area (Fig.2).

In addition to the locations of all currently known thick-billed parrot nest sites, the *nestR* method indicated the presence of a previously undetected active nest site near the city of Madera, Chihuahua. The location of this new nest site was confirmed via ground truthing to be 99 m from its predicted location. The nest was used during the 2022 breeding season for 66 days by a parrot that was telemetered in 2021 at a nearby Madera nest site 5.7 km away (Fig.3). Individual and mean population-level coefficients of habitat selection derived from the Habitat-Selection Function both indicated a positive spatial association of thick-billed parrots with landcover areas characterized by high elevation, taller trees, and trees with wider trunk diameters (β > 0.3, *p* <0.05). Mean elevation for the nesting AKDE home range was 2004 m (min = 577, max = 2665). Mean elevation for the overwintering range was lower with a greater range at 1508 m (min = 67, max = 3129). Mean maximum tree height within the total AKDE home range was 6.7 m (± 4.8), mean maximum tree diameter was 15.3 cm (± 8.1 SD), and the mean tree density was 574 trees/ha (± 320.4 SD). Association of the tracked parrots with forest species type was only weakly positive, and there was no significant association of parrots with landscapes of increasing tree density (Fig.4, Table 1.).

**Table 1.** Population-level mean coefficients for environmental covariates influencing the spatial associations of the tracked thick-billed parrots derived from the independent random slopes model (mixed model approach of Muff et al. 2020)

| | $\beta$ Estimate | Std. Error | z value | P |
| --- | --- | --- | --- | --- |
| Elevation | 0.353 | 0.144 | 2.45 | **0.01** |
| Forest type | -0.049 | 0.023 | -2.12 | **0.03** |
| Tree diameter | 0.621 | 0.076 | 8.16 | **<0.01** |
| Tree density | -0.056 | 0.091 | -0.62 | 0.54 |
| Tree height | 0.689 | 0.108 | 6.39 | **<0.01** |

**Fig 4.**
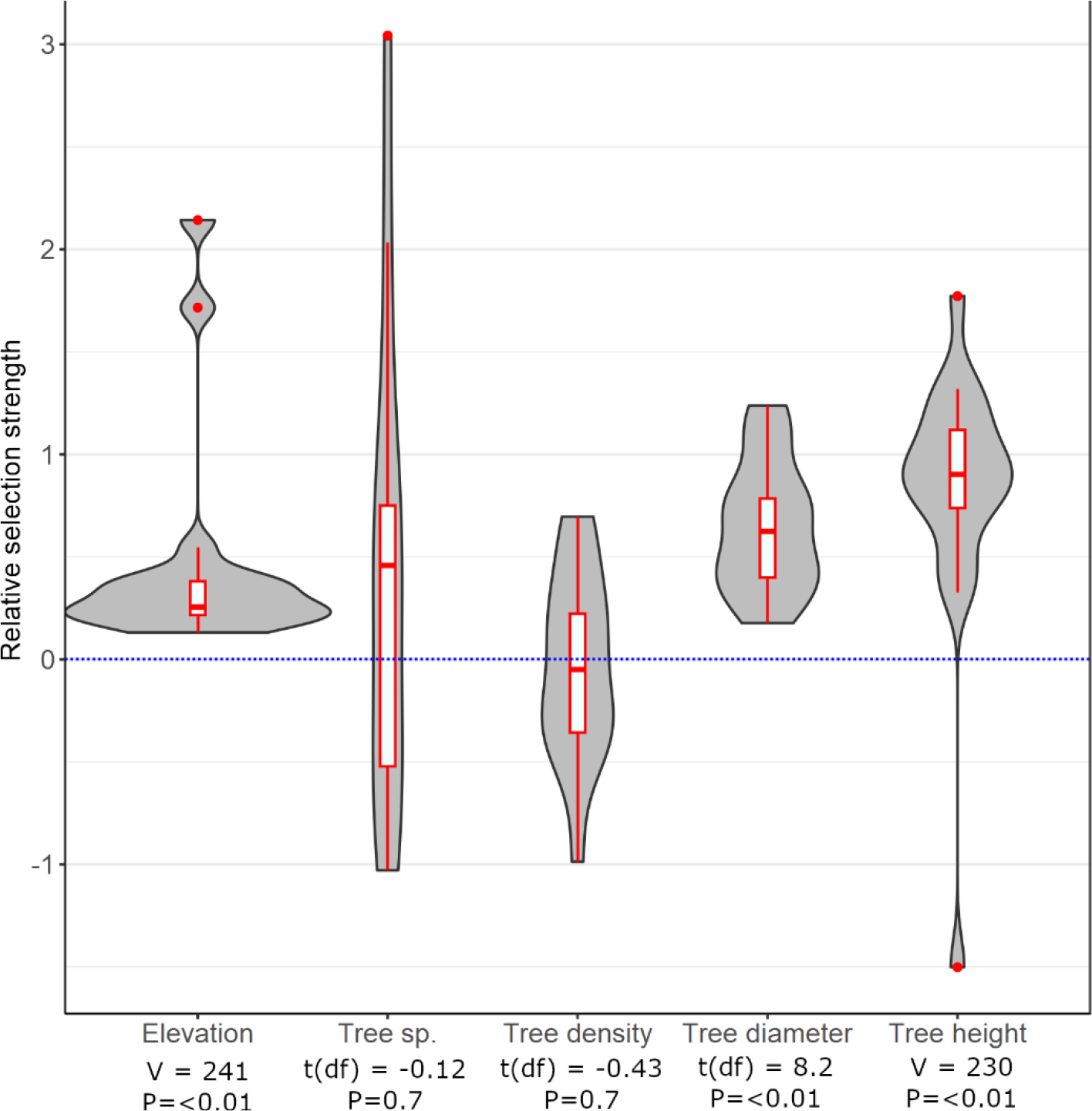
Mean individual regression coefficients with SE in fitted habitat-selection functions for landscape classes within the ranges of the tracked parrots (two-step method of Fieberg et al. 2021), enabling exploration of among-animal variability (indicated by the results of one-sample t-tests or Wilcoxon tests).

We detected variations in seasonal home range sizes for the tracked thick-billed parrots, as well as matching seasonal variations in the area of parrot home range overlap. The mean breeding site AKDE home range area was much smaller (7,468 km^2^, 3,585 -13,640 CI) than the mean overwintering home range area (19,993 km^2^, 13,206 - 28,845 CI), and this significant difference was confirmed via paired *t*-test (*t* = -3.08, df = 15, *p* = 0.008). The mean breeding site overlap based on the Bhattacharyya coefficient was also much smaller (0.38, 0.32 - 0.34 CI) than the mean overwintering overlap estimate (0.73, 0.68 - 0.78 CI), and this difference was confirmed via paired t-test (t = -7.94, df = 19, *p* = <0.001). The mean Kriged occurrence distribution estimate of parrot movements provided a population-level estimate of the dimensions of combined migration routes between the two seasonal ranges (Fig.5). This model also identified the locations of five previously unknown stopover sites on remote forested high-elevation plateaus and ridgelines along the Sierra Madre Occidental ranges between the towns of Guachochi and Choix (Fig.5, Appendix 4). The tracked parrots tended to depart their seasonal habitats together within close social groups around the same time periods. We observed six groups comprised of 3 – 7 parrots each that were highly spatially associated (Bhattacharyya coefficient overlap values 0.7 - 0.97) and departed either the nesting or overwintering habitats within 3 - 10 days of each other. Most of the habitat within thick-billed parrot home ranges currently lack formal protection. As of 2023, only 19.6 % of the total thick-billed parrot nesting home range area, 8.5 % of the total overwintering home range area, and 6.4 % of the total area of migratory stopover sites that we calculated are covered by regulatory protections, such as a reserve or natural protected area designation (Appendix 5.).

**Fig 5.**
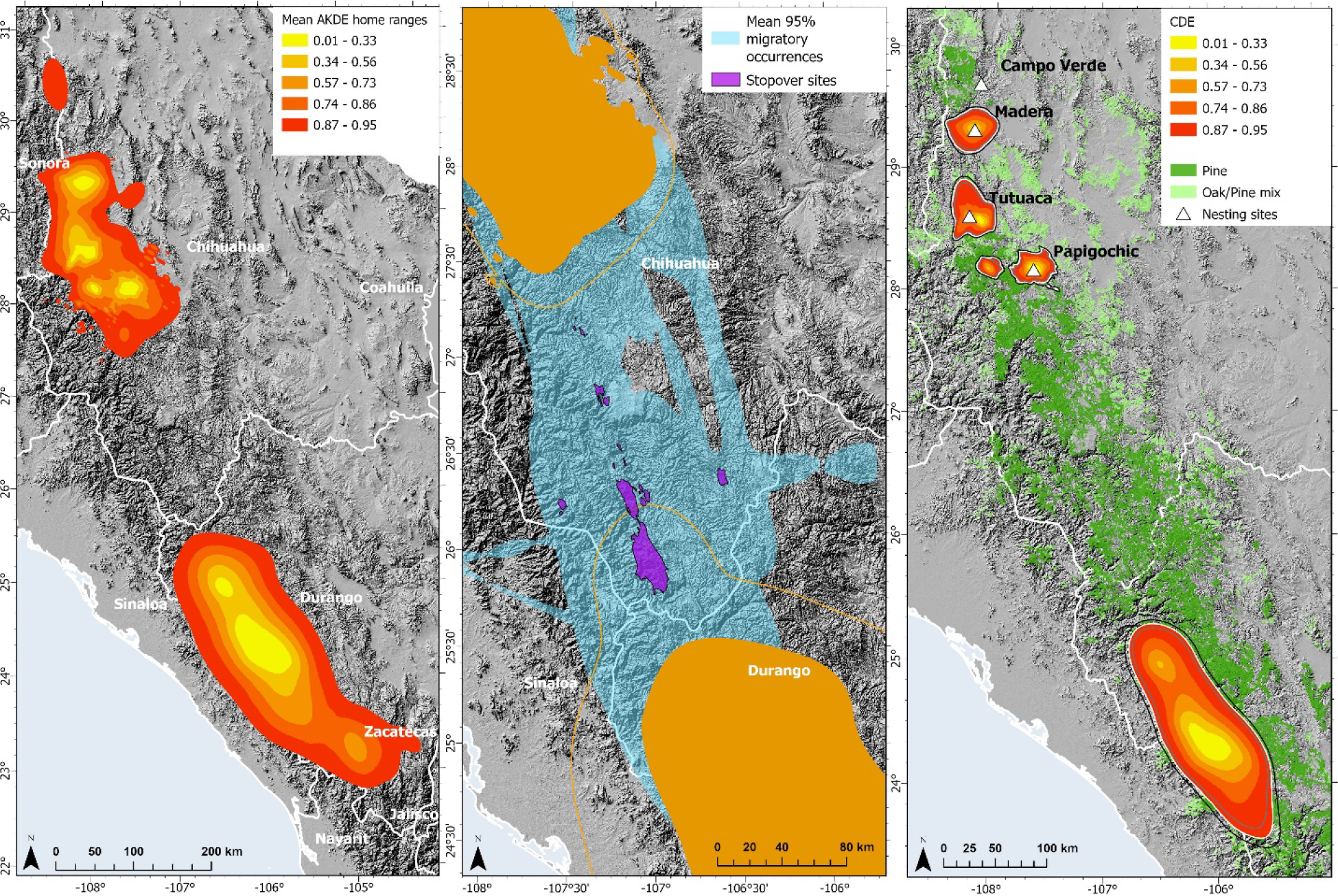
Left: Mean AKDE home ranges with CIs for thick-billed parrot locations recorded at the northern breeding sites and southern overwintering sites. Center: Mean migratory route area defined via 95% Kriged occurrence distribution estimate (blue polygon) with five stopover sites defined by the 25% occurrence contour (purple polygons). Orange polygons indicate seasonal home range areas. Right: Conditional distribution of encounters for parrots at the breeding and overwintering sites indicating a higher likelihood of encounters concentrated around the nesting locations.

## Discussion

Our study elucidates the locations of hitherto undefined overwintering home ranges for thick-billed parrots, the spatial dimensions of their migratory routes, and the locations of key migratory stopover sites. We also characterized spatiotemporal patterns of thick-billed parrot migratory behaviors and social interactions and identified their spatial associations with landscape types. All tracked parrots left their breeding areas at a certain time each year and returned at another time, and the timing, directions and distances of their migrations were also relatively constant and predictable from tracking year to year. These spatial behaviors may be evidence that thick-billed parrots are obligate migrators (Newton 2012), although additional tracking and population monitoring will be required to confirm.

Investigations of similar migratory species show that identification of landscape locations and variables important for supporting population persistence provide invaluable information for targeted conservation action, such as habitat management strategies and establishment of protected areas along the entire migratory path (Runge et al. 2015, Schuster et al. 2019).

Most parrot species can switch dietary items in response to fluctuating food availability. However, Psittaciformes with specialized diets such as thick-billed parrots tend to consume foods that are available throughout the year and are likely to be highly mobile to track resource patches and increase foraging efficiency (see review by Renton et al. 2015). All the parrots that we telemetered undertook migrations and exhibited range shifting behavior. Although there was high intraspecific variation in spatial behaviors, most of the tracked parrots departed their seasonal ranges within a few weeks of each other and traveled along similar migratory routes in close social groups. Migrating parrots followed the high-elevation forests along the ridgeline of the Sierra Madre Occidental ranges in mostly linear flight paths. Most of the tracked parrots also spent some time at stopover sites during their migrations, which underscores their importance as key components of thick-billed parrot habitat. Boundaries of parrot space-use patterns were smaller at the northern breeding ranges than the southern overwintering ranges, which likely reflects how mating, nesting and chick-rearing behaviors constrain parrot movements. For example, breeding pairs are limited in how far they can range each day by the need to regularly return to their nest and feed their developing chicks. Alternatively, smaller home ranges at the breeding sites may reflect greater competition for resources during the reproductive season. Food resources may also be more dispersed around overwintering ranges compared with those around nesting sites, which would require larger and longer foraging trips.

Positive spatial association of the tracked parrots to landscapes containing tall trees with wide diameters accords with prior suggestions that thick-billed parrot populations require old growth forests comprised of mature pine trees that provide nesting cavities and pine seed food resources (Lanning and Shiflett 1983, Snyder et al. 1995). This finding is also consistent with the growing body of evidence that old-growth forests are crucial for the conservation of cavity-nesting Psittaciformes and other avian species (e.g. Lammertink et al. 1996, Webb et al. 2012, Manning et al. 2013, de la Parra-Martínez et al. 2015). Absence of parrot spatial association with regions of increasing tree density may be because the taller and broader trees that parrots prefer tended to occur within less dense forests (see Zeide 1995), or our result may be an artifact of insufficient model rigor or mismatched granularity between this landscape layer and the parrot movement data. Although there was some variability, e.g., when flying across low-level areas between core habitats, our study confirms the preference of thick-billed parrots for forested landscapes of high elevation. We can also infer from our model of spatial association with forest species type that the parrots use habitats that are either predominantly conifer or conifer-oak mixes rather than habitats that are only conifer dominated. This behavior may reflect selection for a broader niche of tree species used for nesting cavities or to patterns in food availability. Although the thick-billed parrot primarily consumes the seeds of conifers, the species has also been reported feeding on acorns and the buds of various trees (Snyder et al. 1995). Hence, our results may reflect greater foraging flexibility than was presupposed, or a response to reduced pine seed availability due to drought or wildfires.

Future studies of thick-billed parrot spatial ecology should explore thick-billed parrot decision rules and phenotypic responses to environmental conditions that trigger migration events. Seasonal migrations may be prompted by specific temperature cues and/or daylight thresholds. The pine species that comprise the bulk of the parrot’s food show strong differences in the regularity of fruiting and in their seasonal availability of seeds (Snyder et al. 1995). Hence, parrot spatial behaviors may also be driven by this variability in food resource availability, especially during drought conditions. Future investigations should also derive robust estimates of habitat carrying capacity to inform models for defining the amount of habitat required to support parrot populations and enable recovery to self-sustaining levels. Future studies should also define the functional role of thick-billed parrots in the processes of pollination, seed dispersal, and seed predation and their broader role in maintaining the ecological health of the Sierra Madre Occidental bioregion. Psittaciformes are among the most endangered birds with the main threats to wild Neotropical parrot populations related to human activities, principally agro-industry, farming and grazing (see review by Berkunsky et al. 2017). Suitable tree cavities are an essential component of thick-billed parrot breeding habitat. Availability of nesting cavities in mature trees and snags in the Sierra Madre Occidental has dramatically decreased in recent decades due to destructive clear-cutting and high-intensity timbering of remnant old-growth primary forests, which highlights the need to: (1) Conserve remaining old growth; (2) allow secondary growth to mature, and; (3) experiment with installation of artificial cavities similar to that of Tarazona-Tubens et al. (2022) (e.g. to better understand the microclimate and microhabitat features that promote nesting success).

Habitat restoration and protection is one of the most common management strategies to improve the conservation outlook for Neotropical parrot populations (Berkunsky et al. 2017). The lack of formal protection currently afforded to thick-billed parrot habitats within their range is likely a serious impediment to the protection and recovery of their remaining populations. Existing networks of protected sites may also not be adequate under future scenarios of environmental change. For example, important foraging and nesting trees for parrots that are already threatened by anthropogenic disturbance may be further impacted by increasing intensity and frequency of forest fire events and pine beetle outbreaks exacerbated by climate change (Sáenz-Romero et al. 2010). Plasticity in diet and foraging strategy influences the extent to which parrots can respond to anthropogenic pressures of global change (Renton et al. 2015). The dependence of thick-billed parrots on seeds from high-elevation pine species may severely restrict their ability to adapt in time to increasingly fluctuating resource availability (e.g. via habitat shifting). Thick-billed parrot functional responses to foraging resources may also become mismatched if the environmental cues prompting seasonal migrations become disassociated from the seasonality of food resources (see Brown et al. 2016). We have observed drastic reductions in parrot breeding success during drought periods that impact pine tree seed productivity (unpublished data). Precipitation scenarios in recent climate change assessments project large probabilities of decreases in rainfall over northwestern Mexico (Magaña et al. 2012), which will likely have major long-term consequences for thick-billed parrot reproductive output.

### Management implications

Our findings should be integrated into current and future landscape and natural resource management plans to enhance the sustainable use of the Sierra Madre Occidental bioregion and improve the conservation outlook for thick-billed parrot populations across their annual life cycle. Somewhat counterintuitively, we found that parrots need larger protected areas to cover their non-breeding range. Resource management for foraging resources may be even more important in their breeding season range because longer foraging movements are constrained by nesting, so increased protection measures should focus on preserving key pine seed foraging resources that are local to nest sites. Specifically, we recommend that formal habitat protections be reviewed and expanded to cover key nesting sites in western Chihuahua/eastern Sonora, as well as overwintering habitat in western Durango/eastern Sinaloa. The importance of acquiring sufficient energetic resources along the migratory path for birds that undertake long-distance movements is well established (Hutto 1998, Weber et al. 1999). Hence, migratory stopover sites in the mountains of southern Chihuahua should also receive increased protection.

Successful stabilization and recovery of wild populations of thick-billed parrots and protection of the old-growth forest that support them will not be possible without sustained engagement with and support from the local stakeholders within parrot ranges, including ejido communities and forestry industry groups (Lamberski and Healy 2002). Improved understanding of the timing of parrot seasonal migrations presented here may inform regulatory decisions and community outreach initiatives to minimize disturbance at key seasonal, stopover, and migratory pathway locations at specific times of the parrots’ annual life cycle. Our findings will inform future considerations of reintroductions and translocations of thick-billed parrot populations into their former U.S. ranges and provide a reference to identify suitable habitat within the U.S. to set potential future restoration targets. More broadly, our findings contribute to the rapidly growing application of movement ecology to conservation. Indeed, more than one-third of all studies in this discipline make explicit conservation applications, and the uptake by government agencies (e.g., in recovery plans) is a remarkable 60%, indicating that findings are readily understood and translated into conservation action (Fraser et al. 2018). Future work should ensure that these findings can be applied to the conservation of thick-billed parrots to the greatest extent possible.

## Ethics statement

Protocols for handling and telemetering of wild thick-billed parrots were reviewed and approved by the Subsecretaría de Gestión Para la Protección Ambiental, Dirección General De Vida Silvestre, Mexico Federal Government (Undersecretariat of Management for Environmental Protection, General Directorate of Wildlife) (Project # 09/K6-0820/03/l9) as well as the San Diego Zoo Wildlife Alliance Institutional Animal Use and Care Committee (Project #s 18-019 and 15-006).

## Data accessibility

Argos telemetry data used in this study are available with acceptance of license terms from the movebank online data repository (Movebank ID# 2842491348).

## Acknowledgements

This research was supported with the assistance of the Comisión Nacional de Áreas Naturales Protegidas (CONANP) through their Programa para la Protección y Restauración de Ecosistemas y Especies en Riesgo (PROREST 2019). Project support was also provided by the Arizona Game and Fish Department (AZGFD), Organización Vida Silvestre A.C. (OVIS), the Universidad Autónoma de Nuevo León (UANL), and the San Diego Zoo Wildlife Alliance. The authors wish to acknowledge and thank the following individuals and organizations for their support for this project: World Parrot Trust, Ellen Browning Scripps Foundation, William E. Cole Family Foundation, Sacramento Zoo, U.S. Fish & Wildlife Service, Nacho Vilchis, Yvette Kemp, Edwin Juarez, Angie Olvera, Francisco Puente, Mateo Rangel, Jesús Márquez, Abel Guerrero, Horacio Barcenas, the Ejidos: El Largo, J. Rojo Gómez and Tutuaca. All of the authors of this study have no conflict of interest to declare.

## Notes

### Competing Interest Statement

The authors have declared no competing interest.

### Summary of Updates

We have contextualized the population status of thick-billed parrots with direct reference to IUCN RedList criteria. We have highlighted the importance of our work in relation to parrot conservation in general, and contrasted our findings to what is known (or unknown) about other parrot species. And where necessary, we have provided additional clarifications to our terminology and spatial modeling processes and outputs.

