## Supplementary for "Spatial behaviors and seasonal habitat use of the increasingly endangered thick-billed parrot (*Rhynchopsitta pachyrhyncha*)"

**Appendix 1:** Characteristics of the thick-billed parrots that were captured and telemetered in the northern Sierra Madre Occidental mountain ranges of Mexico between 2019 – 2022.

| Bird ID | Date tagged | Weight (g) | Wing width (mm) | Age | Site released |
|---|---|---|---|---|---|
| TBP1 | 19-Sep-19 | 365 | 245 | 55 days | Madera |
| TBP2 | 29-Sep-19 | 389 | 230 | 58 days | Madera |
| TBP3 | 24-Sep-19 | 397.8 | 264 | Adult | Tutuaca |
| TBP4 | 21-Sep-19 | 356 | 238 | 55 days | Papigochic |
| TBP5 | 21-Sep-19 | 382 | 268 | Adult | Papigochic |
| TBP6 | 20-Sep-19 | 396 | 256 | Adult | Madera |
| TBP7 | 2-Oct-19 | 330 | 242 | 57 days | Papigochic |
| TBP8 | 1-Oct-19 | 396 | 230 | 53 days | Campo Verde |
| TBP9 | 24-Sep-19 | 354.2 | 255 | Adult | Tutuaca |
| TBP10 | 2-Oct-19 | 371 | 226 | 55 days | Papigochic |
| TBP11 | 24-Sep-20 | 335 | 258 | Adult | Janos |
| TBP12 | 26-Sep-20 | 364.9 | 257 | Adult | Madera |
| TBP13 | 26-Sep-20 | 332.2 | 225 | 56 days | Madera |
| TBP14 | 30-Sep-20 | 350 | 201 | 52 days | Tutuaca |
| TBP15 | 1-Oct-20 | 340.2 | 250 | Adult | Tutuaca |
| TBP16 | 1-Oct-20 | 355.9 | 256 | Adult | Tutuaca |
| TBP17 | 1-Oct-20 | 337.4 | 230 | 57 days | Tutuaca |
| TBP18 | 4-Oct-20 | 373.5 | 265 | Adult | Papigochic |
| TBP19 | 4-Oct-20 | 403.7 | 250 | Adult | Papigochic |
| TBP20 | 4-Oct-20 | 380 | 204 | 54 days | Papigochic |
| TBP11 | 30-Sep-21 | 327.1 | 251 | Adult | Janos |
| TBP21 | 3-Oct-21 | 317.8 | 234 | 56-60 Days | Campo Verde |
| TBP22 | 6-Oct-21 | 352.01 | 249 | Adult | Tutuaca |
| TBP23 | 6-Oct-21 | 336.2 | 257 | Adult | Tutuaca |
| TBP24 | 8-Oct-21 | 401 | 255 | Adult | Madera |
| TBP25 | 8-Oct-21 | 365.2 | 260 | Adult | Madera |
| TBP26 | 8-Oct-21 | 363.5 | 204 | 54 Days | Madera |
| TBP27 | 10-Oct-21 | 352.8 | 263 | Adult | Papigochic |
| TBP28 | 10-Oct-21 | 360.9 | 270 | Adult | Papigochic |
| TBP29 | 10-Oct-21 | 371.6 | 225 | 55 Days | Papigochic |
| TBP30 | 10-Sep-22 | 379.9 | 260 | Adult | Campo verde |
| TBP31 | 15-Sep-22 | 393.8 | 263 | Adult | Janos |
| TBP32 | 19-Sep-22 | 373.7 | 265 | Adult | Tutuaca |
| TBP33 | 21-Sep-22 | 291.8 | 238 | 57 days | Madera |

**Appendix 2:** Range shift model outputs estimated using the likelihood method for the Ornstein-Uhlenbeck position process. Range shifts were confirmed by testing the l.r.t with 4 d.f against no migration. Stopover ranges were confirmed by testing whether the AIC of a model fitted using a three-range model was lower than the AIC of a two-range model. **Or** = general orientation of migratory movement towards the south (S) away from the breeding sites or north (N) indicating a return migration. **Cen** = number of centroids fitted in the range model. Leave and Arrive dates are month + tracking year. ╡$t_k$ = duration of range transition in days. ***D*** = the distance between the centroids of the respective ranges in km. Increasing range shift index (**RSI**) values indicate longer migrations. Significance symbols (***, **, -) represent *P*-values below 0·001, 0·01, and above 0·05 respectively.

| ID | *n* | Or | Cen | l.r.t | Shift *P* | Stop *P* | Leave | Arrive | ╡$t_k$ | *D* | RSI | 95% CI |
|---|---|---|---|---|---|---|---|---|---|---|---|---|
| TBP2 | 288 | S | 3 | 233.7 | *** | Y *** | Oct-19 | Nov-19 | 29 | 576 | 3.4 | 3.2-3.6 |
| TBP3 | 313 | S | 3 | 187 | *** | Y *** | Oct-19 | Nov-19 | 20 | 330 | 2.8 | 2.6-3.0 |
| TBP3 | 334 | N | 3 | 68.9 | *** | Y * | Apr-20 | May-20 | 28 | 188 | 2.2 | 2.1-2.3 |
| TBP4 | 319 | S | 3 | 136.4 | *** | Y *** | Oct-19 | Nov-19 | 46 | 417 | 1.6 | 1.5-1.7 |
| TBP4 | 156 | N | 3 | 38.2 | *** | Y *** | Apr-20 | May-20 | 17 | 299 | 1.9 | 1.7-2.0 |
| TBP4 | 212 | S | 2 | 41 | *** | N - | Aug-20 | Aug-20 | 9 | 185 | 1.4 | 1.3-1.5 |
| TBP5 | 376 | S | 3 | 31.8 | *** | N - | Oct-19 | Dec-20 | 70 | 475 | 2.0 | 1.9-2.1 |
| TBP6 | 321 | S | 2 | 41.1 | *** | N - | Oct-19 | Nov-19 | 36 | 517 | 2.5 | 2.4-2.6 |
| TBP6 | 114 | N | 3 | 63.7 | *** | Y *** | Apr-20 | May-20 | 15 | 498 | 4.7 | 4.4-5.2 |
| TBP6 | 162 | S | 2 | 82.6 | *** | N - | Oct-20 | Oct-20 | 8 | 714 | 3.7 | 3.5-4.0 |
| TBP7 | 297 | S | 3 | 14.2 | *** | Y ** | Oct-19 | Jan-20 | 74 | 564 | 2.8 | 2.6-3.0 |
| TBP8 | 137 | S | 2 | 67.3 | *** | Y * | Oct-19 | Oct-19 | 11 | 173 | 3.3 | 3.0-3.6 |
| TBP9 | 313 | S | 2 | 188.2 | *** | N - | Oct-19 | Oct-19 | 4 | 529 | 4 | 3.8-4.2 |
| TBP9 | 291 | N | 2 | 70.8 | *** | N - | Apr-20 | Apr-20 | 6 | 412 | 3.7 | 3.5-4.0 |
| TBP9 | 278 | S | 3 | 124 | *** | Y *** | Oct-20 | Dec-20 | 73 | 317 | 1.0 | 0.9-1.1 |
| TBP10 | 201 | S | 3 | 151.2 | *** | Y *** | Oct-19 | Nov-19 | 28 | 421 | 2.7 | 2.5-2.9 |
| TBP12 | 309 | S | 2 | 182.2 | *** | N - | Oct-20 | Oct-20 | 12 | 765 | 4.1 | 3.9-4.3 |
| TBP12 | 570 | N | 3 | 149 | *** | Y *** | Apr-21 | Jun-21 | 45 | 744 | 6.1 | 5.9-6.4 |
| TBP12 | 360 | S | 3 | 20 | *** | Y *** | Oct-21 | Nov-21 | 41 | 569 | 4.1 | 3.9-4.4 |
| TBP12 | 270 | N | 2 | 63.5 | *** | N - | Apr-22 | Apr-22 | 19 | 485 | 1.9 | 1.8-2.0 |
| TBP14 | 301 | S | 2 | 1344.8 | *** | N - | Oct-20 | Oct-20 | 4 | 216 | 12.3 | 11.7-13.0 |
| TBP15 | 300 | S | 2 | 177.5 | *** | N - | Oct-20 | Oct-20 | 11 | 506 | 2.6 | 2.5-2.8 |
| TBP16 | 120 | S | 3 | 139 | *** | N - | Oct-20 | Oct-20 | 3 | 550 | 3.7 | 3.4-4.1 |
| TBP16 | 471 | N | 3 | 173 | *** | Y *** | Dec-20 | Jun-21 | 181 | 305 | 2.1 | 2.0-2.2 |
| TBP18 | 293 | S | 3 | 58.3 | *** | Y ** | Oct-20 | Nov-20 | 43 | 284 | 1.1 | 1.1-1.2 |
| TBP17 | 126 | S | 2 | 32.3 | *** | N - | Oct-20 | Oct-20 | 3 | 470 | 3.1 | 2.7-3.6 |
| TBP18 | 156 | N | 3 | 66.7 | *** | Y ** | May-21 | May-21 | 5 | 359 | 2.9 | 2.7-3.2 |
| TBP18 | 360 | S | 2 | 114.8 | *** | N - | Nov-21 | Dec-21 | 15 | 449 | 4.4 | 4.2-4.6 |
| TBP22 | 298 | S | 3 | 110.1 | *** | Y *** | Oct-21 | Nov-21 | 42 | 537 | 2.5 | 2.4-2.7 |
| TBP22 | 190 | N | 3 | 127 | *** | Y *** | Apr-22 | Apr-22 | 5 | 596 | 2.1 | 1.9-2.2 |
| TBP23 | 345 | S | 2 | 315 | *** | Y *** | Oct-21 | Nov-21 | 40 | 480 | 4 | 3.8-4.3 |
| TBP23 | 112 | N | 3 | 243 | *** | Y *** | Apr-22 | Apr-22 | 8 | 403 | 4.2 | 3.9-4.7 |

| | | | | | | | | | | | | |
|---|---|---|---|---|---|---|---|---|---|---|---|---|
| TBP23 | 271 | S | 2 | 28.2 | *** | Y *** | Sep-22 | Oct-22 | 48 | 352 | 1.7 | 1.6-1.8 |
| TBP24 | 285 | S | 2 | 43.1 | *** | N - | Oct-21 | Nov-21 | 31 | 553 | 2.4 | 2.3-2.6 |
| TBP24 | 192 | N | 2 | 143 | *** | N - | Apr-22 | Apr-22 | 6 | 399 | 6.8 | 6.3-7.7 |
| TBP24 | 268 | S | 2 | 55.5 | *** | Y *** | Sep-22 | Oct-22 | 25 | 498 | 2.9 | 2.7-3.1 |
| TBP25 | 286 | S | 2 | 49.7 | *** | N - | Oct-21 | Nov-21 | 28 | 548 | 2.9 | 2.7-3.0 |
| TBP28 | 281 | S | 2 | 105.5 | *** | N - | Nov-21 | Dec-21 | 10 | 428 | 3.5 | 3.4-3.7 |
| TBP28 | 337 | N | 2 | 69.5 | *** | N - | Apr-22 | May-22 | 13 | 342 | 2.3 | 2.2-2.4 |
| TBP29 | 281 | S | 3 | 132.2 | *** | Y *** | Nov-21 | Nov-21 | 21 | 410 | 3.1 | 3.0-3.3 |
| TBP29 | 121 | N | 3 | 83.7 | *** | Y *** | May-22 | May-22 | 4 | 458 | 2.8 | 2.6-3.1 |
| TBP29 | 107 | S | 2 | 192 | *** | N - | Sep-22 | Sep-22 | 5 | 226 | 5.1 | 4.8-5.5 |
| TBP31 | 237 | S | 2 | 389.7 | *** | N - | Oct-22 | Oct-22 | 4 | 565 | 2.2 | 2.1-2.4 |
| TBP32 | 230 | S | 2 | 64.7 | *** | N - | Sep-22 | Oct-22 | 19 | 464 | 3.2 | 3.0-3.4 |

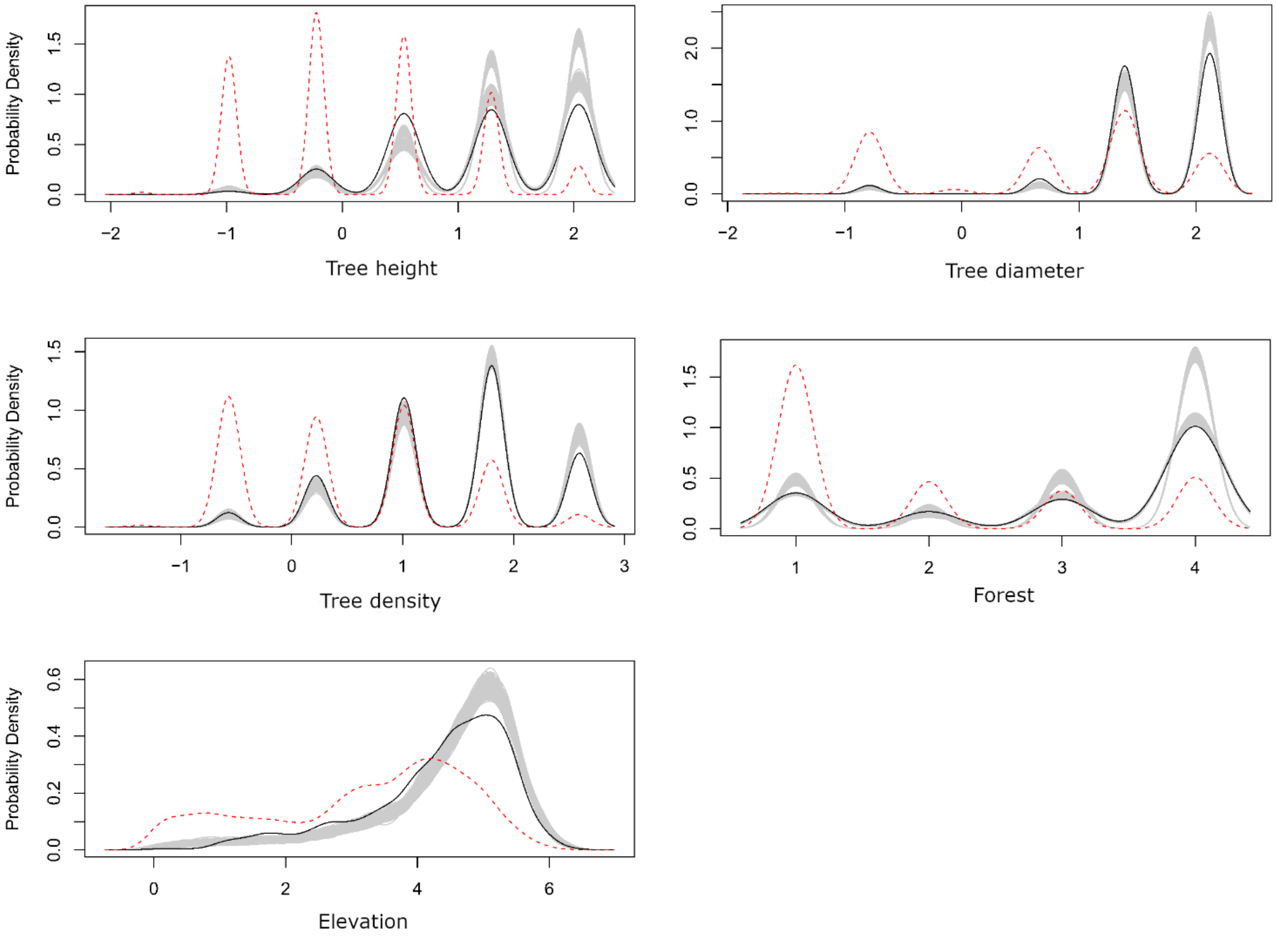


**Appendix 3.**

Used-habitat calibration plots for habitat-selection models fit to the thick-billed parrot telemetry data. Scaled covariates in the models measured coverage of tree height, tree diameter, tree density, forest quality based on dominant tree species, and elevation. Panels depict the distribution of available and used locations in the test data set (red dashed and solid black lines, respectively), with a 95% simulation envelope for these distributions given by the gray bands. A model is considered well-calibrated if the observed distributions (solid black lines) fall within the simulation envelopes (Fieberg et al. 2018).

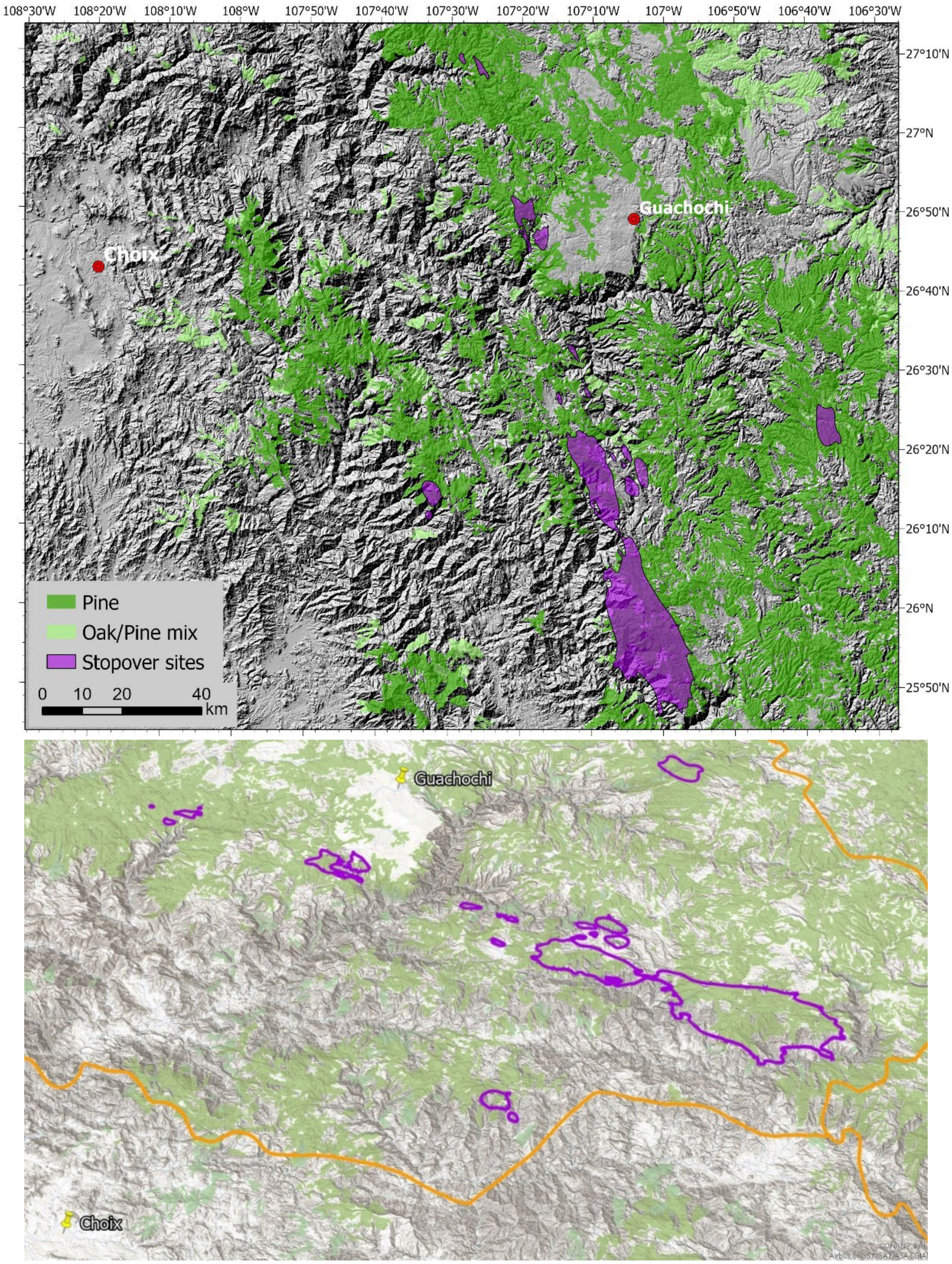


**Appendix 4.**

2D and 3D closeups of the locations of five previosuly unknown stopover sites used by the tracked parrots during their north/south seasonal migrations. Stopover polygons were derived from the 25% contours of the Kriged occurrence distribution estimate of parrot movements These stopover sites were located on high-elevation forest plateaus and ridgelines between the towns of Choix (108°19'33"W 26°42'34"N ) and Guachochi (107°4'10"W 26°49'13"N).

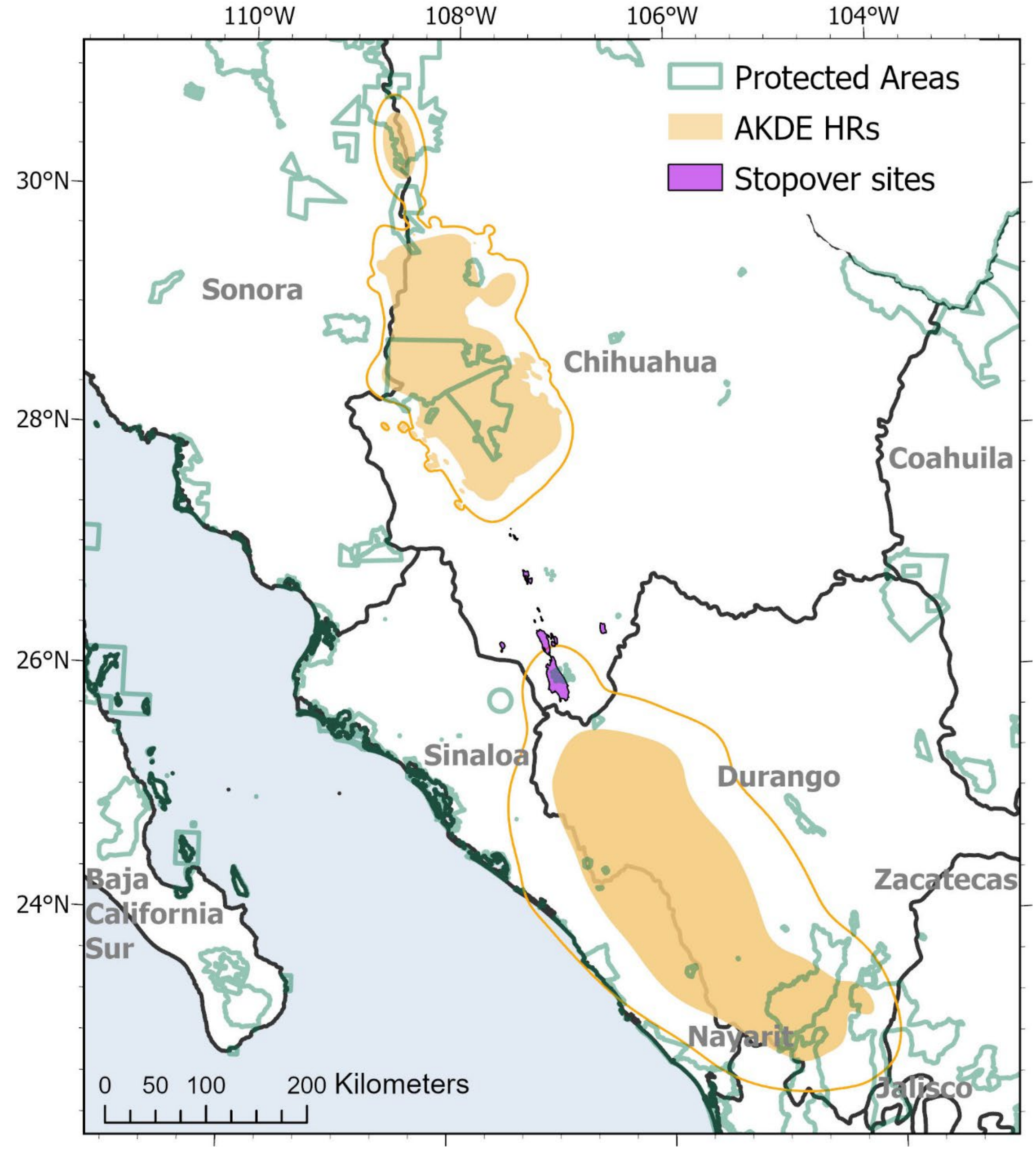


**Appendix 5.**

Northern breeding and southern overwintering AKDE home ranges for thick-billed parrots (orange polygons) overlaid with current protected area zones (green polygons) designated by the State and Federal Government of Mexico. As of 2023, only 19.6 % of the total thick-billed parrot nesting home range area, 8.5 % of the total overwintering home range area, and 6.4 % of the total area of migratory stopover sites that we calculated are covered by formal regulatory protections.
